# Disconnect between signalling potency and *in vivo* efficacy of pharmacokinetically optimised biased glucagon-like peptide-1 receptor agonists

**DOI:** 10.1101/855874

**Authors:** Maria Lucey, Philip Pickford, James Minnion, Jan Ungewiss, Katja Schoeneberg, Guy A Rutter, Stephen R Bloom, Alejandra Tomas, Ben Jones

## Abstract

**Objective:** To determine how pharmacokinetically advantageous acylation impacts on glucagon-like peptide-1 receptor (GLP-1R) signal bias, trafficking, anti-hyperglycaemic efficacy and appetite suppression.

**Methods:** *In vitro* signalling responses were measured using biochemical and biosensor assays. GLP-1 receptor trafficking was determined by confocal microscopy and diffusion-enhanced resonance energy transfer. Pharmacokinetics, glucoregulatory effects and appetite suppression were measured in acute, sub-chronic and chronic settings in mice.

**Results:** A C-terminally acylated ligand, exendin-phe1-C16, was identified with undetectable β-arrestin recruitment and GLP-1R internalisation. Depending on the cellular system used, this molecule was up to 1000-fold less potent than the comparator exendin-asp-3-C16 for cyclic AMP signalling, yet was considerably more effective *in vivo*, particularly for glucose regulation.

**Conclusions:** C-terminal acylation of biased GLP-1R agonists increases their degree of signal bias in favour of cAMP production and improves their therapeutic potential.

## 1 Introduction

Glucagon-like peptide-1 receptor agonists (GLP-1RAs) are effective agents for the treatment of type 2 diabetes (T2D) and obesity (1). Their therapeutic effects derive mainly from potentiation of glucose-stimulated insulin secretion and suppression of appetite leading to weight loss, respectively mediated by GLP-1Rs expressed in pancreatic beta cells and anorectic neurons. GLP-1RAs are known to improve renal (2) and cardiovascular outcomes (3) in T2D, and reduce mortality (4).

The predominant intracellular signalling intermediate that couples GLP-1R activation to its downstream effects is cyclic adenosine monophosphate (cAMP) (5, 6). However, an updated view of GLP-1R pharmacology highlights the roles of membrane trafficking (7, 8) and additional effector proteins such as the β-arrestins (9, 10) in the control of amplitude, duration and subcellular localisation of signalling events to regulate insulin secretion, particularly in the pharmacological setting. Although all clinically approved GLP-1RAs show broadly similar signalling and trafficking characteristics to the endogenous ligand, GLP-1(7-36)NH_2_, these can be dramatically altered via sequence modifications close to the ligand N-terminus, as demonstrated recently using analogues of the GLP-1 homologue peptide exendin-4 (11, 12). Specifically, “biased” GLP-1RAs which retain full efficacy for cAMP production, but reduced β-arrestin recruitment and endocytic uptake, are able to avoid GLP-1R desensitisation and downregulation which ordinarily attenuate glucoregulatory responses *in vivo*.

Studies of orthosteric biased GLP-1R agonism have so far mainly used peptide ligands (11, 12), sometimes featuring non-native amino acid substituents (13–15). Whilst these ligands typically have been engineered for high proteolytic stability, rapid renal elimination (16) means that their half-lives are measured in hours, which is incompatible with a drive to reduce the frequency of injections for patient convenience, comfort and adherence. The leading GLP-1RAs in current clinical usage have been chemically optimised to allow once-weekly dosing in humans through avoidance of renal clearance (1), for example through conjugation of a fatty acid chain to the peptide which promotes reversible binding to albumin and other plasma proteins which are too large to undergo glomerular filtration. However, these approved compounds show broadly comparable signalling characteristics, and biased GLP-1R agonism has not yet been studied using pharmacokinetically optimised agents.

In this study, we investigate the impact of acylating two oppositely biased GLP-1R agonists, exendin-phe1 and exendin-asp3, both with identical amino acid sequences to exendin-4 except for at the first or third N-terminal amino acids, respectively (12). We found that the introduction of a C-terminal fatty acid chain exaggerated the degree of bias at the expense of a reduced overall signalling potency for exendin-phe1-C16. However, when tested *in vivo*, despite up to ∼1000-fold reduction in signalling potency, exendin-phe1-C16 outperformed exendin-asp3-C16 for control of blood glucose over 72 hours after a single dose, as well as providing greater glucose control and weight loss with repeated administration.

## 2 Materials and methods

### 2.1 Peptides

Peptides were obtained from Wuxi AppTec Co., Ltd., and were of >90% purity.

### 2.2 Cell culture

HEK293 cells stably expressing human SNAP-GLP-1R (17) were maintained in DMEM with 10% foetal bovine serum (FBS), 1% penicillin/streptomycin and G418 (1 mg/ml). HEK293T cells were maintained similarly but without G418. PathHunter CHO-K1-βarr2-EA-GLP-1R cells (DiscoverX) were maintained in F12 medium with 10% FBS, 1% penicillin/streptomycin, G418 (1 mg/ml) and hygromycin (250 µg/ml). INS-1 832/3 cells (a gift from Prof. Christopher Newgard, Duke University) and MIN6B1 cells (a gift from Prof Philippe Halban, University of Geneva) were maintained as previously described (12)

### 2.3 GLP-1R binding affinity measurement

HEK293-SNAP-GLP-1R cells were labelled in suspension with SNAP-Lumi4-Tb (Cisbio, 40 nM) for 1 hour at room temperature in complete medium. After washing and resuspension in Hank’s buffered salt solution (HBSS) containing 0.1% bovine serum albumin (BSA) and metabolic inhibitors to prevent GLP-1R internalisation [20 mmol/L 2-deoxygucose and 10 mmol/L NaN_3_ (18)], cells were treated with 10 nM exendin(9-39)-FITC in competition with a range of concentrations of unlabelled peptide for 24 hours at 4°C before measurement of binding by TR-FRET in a Flexstation 3 plate reader as previously described (17). Binding was quantified as the ratio of fluorescent signal at 520 nm to that at 620 nm, after subtraction of the ratio obtained in the absence of FITC-ligands, and equilibrium binding constants (K_d_) were calculated using Prism 8 (GraphPad Software).

### 2.4 Cyclic AMP responses

Cells were stimulated with agonist for the indicated time period in their respective growth mediums. FBS was used when indicated. Assays were performed at 37°C without phosphodiesterase inhibitors except for with INS-1 832/3 and MIN6B1 cells, where 3-isobutyl-1-methylxanthine (IBMX) was added at 500 µM. At the end of the incubation period, cells were lysed, and cAMP was determined by homogenous time-resolved fluorescence (HTRF, cAMP Dynamic 2 kit, Cisbio).

### 2.5 β-arrestin recruitment

Cells were stimulated for 30 minutes in growth medium without FBS at 37°C. The assay was terminated by addition of PathHunter detection reagents and luminescent signal was read from each well.

### 2.6 NanoBiT assays

The plasmids for mini-G_s_, -G_i_ and -G_q_, each carrying an N-terminal LgBiT tag (19), were a gift from Prof Nevin Lambert, Medical College of Georgia. The plasmid for β-arrestin-2 fused at the N-terminus to LgBiT was obtained from Promega custom assay services (plasmid CS1603B118). The SmBiT was cloned in frame at the C-terminus of the GLP-1R by substitution of the Tango sequence on a FLAG-tagged GLP-1R-Tango expression vector (20), a gift from Dr Bryan Roth, University of North Carolina (Addgene plasmid # 66295). HEK293T cells in 12 well plates were co-transfected for 24 hours using Lipofectamine 2000 and following quantities of DNA: 0.5 µg each of GLP-1R-SmBit and LgBit-mini-G, or 0.05 µg each of GLP-1R-SmBit and LgBit-mini-G with 0.9 µg empty vector DNA (pcDNA3.1). Cells were resuspended in Nano-Glo dilution buffer + fumarazine (Promega) diluted 1:20 and seeded in 96-well half area white plates. Baseline luminescence was measured over 5 minutes using a Flexstation 3 plate reader at 37°C before addition of ligand or vehicle. Responses were normalised to average baseline.

### 2.7 Confocal microscopy

HEK293-SNAP-GLP-1R cells seeded on coverslips were labelled with SNAP-Surface 549 prior to stimulation with indicated agonist (100 nM) for 30 minutes, after which cells were washed in PBS, fixed with 4% paraformaldehyde, mounted in Diamond Prolong mounting medium with DAPI, imaged with a Zeiss LSM780 inverted confocal microscope with a 63x/1.4 numerical aperture oil-immersion objective and analysed in Fiji.

### 2.8 Animal studies

All animal procedures were approved by the British Home Office under the UK Animal (Scientific Procedures) Act 1986 (Project License PB7CFFE7A). Lean male C57Bl/6 mice (8-10 weeks of age, body weight 25-30 g, obtained from Charles River) were maintained at 21-23°C and 12-hour light-dark cycles. *Ad libitum* access to water and normal chow (RM1, Special Diet Services), or diet containing 60% fat to induce obesity and glucose intolerance (D12492, Research Diets) for a minimum of 3 months before experiments, was provided. Mice were housed in groups of four, except for food intake assessments when they were individually caged with one week of acclimatisation prior to experiments.

### 2.9 Pharmacokinetic studies

Mice were injected intraperitoneally (IP) with 100 nmol/kg agonist and blood samples obtained by venesection into lithium heparin-coated capillary tubes. Plasma exendin concentrations were measured using a fluorescent enzyme immunoassay (Phoenix Pharmaceuticals) which recognises the C-terminus of exendin-4. A correction factor was applied to account for reduced recovery of acylated exendin-4 analogues, which was measured to be 50% of that of non-acylated exendin-4 when spiked into mouse plasma.

### 2.10 *ES*calate assay

The *ES*calate assay was performed as described previously (21). Full methodological details are provided in Supplementary Methods.

### 2.11 Glucose tolerance testing

Mice were fasted for 4-5 hours before the test. Bodyweight-adjusted doses of glucose (2 g/kg) were injected IP, with or without agonist prepared within the same injector, as indicated. Blood samples were obtained immediately prior to injection and at 20-minute intervals thereafter, and glucose measured using a handheld glucose meter (GlucoRx).

### 2.12 Food intake

For sub-chronic studies, mice were fasted overnight and diet was returned to the cage immediately after IP injection of agonist, with cumulative intake determined by weighing. For the chronic study, mice were fasted for 4-5 hours during the light phase and diet was returned to the cage immediately after IP injection of agonist at the beginning of the dark phase.

### 2.13 Statistical analyses

Quantitative data were analysed using Prism 8.0 (GraphPad Software). In cell culture experiments, technical replicates were averaged so that each individual experiment was treated as one biological replicate. Dose responses were analysed using 4-parameter logistic fits, with constraints imposed as appropriate. Bias analyses were performed as previously described (12, 22). Statistical comparisons were made by t-test or ANOVA as appropriate, with paired or matched designs used depending on the experimental design. Mean ± standard error of mean (SEM) or individual replicates are displayed throughout. Statistical significance was inferred if p<0.05.

## 3 Results

### 3.1 Impact of C-terminal acylation on *in vitro* pharmacology and trafficking of biased GLP-1RAs

The sequences of exendin-phe1, exendin-asp3, and their acylated equivalents exendin-phe1-C16 and exendin-asp3-C16, are given in Figure 1A. The C16 hexadecanedioic acid moiety was inserted at the peptide C-terminus after a lysine linker. This position was chosen to avoid the potential for the fatty acid to interfere with putative interactions made by the peptide N-terminus with the GLP-1R, which appear to be important for inducing signal bias. The equilibrium affinity of exendin-asp3-C16 was found to be similar to that of exendin-asp3 (Figure 1B, log K_d_ −8.9 ± 0.1 *versus* −9.1 ± 0.1, respectively, p>0.05 by one-way randomised block ANOVA with Sidak’s test), whereas exendin-phe1-C16 showed somewhat higher affinity for the receptor than its non-acylated counterpart (log K_d_ −8.3 ± 0.0 *versus* −7.8 ± 0.0, respectively, p<0.05 by one-way randomised block ANOVA with Sidak’s test).

**Figure 1.**
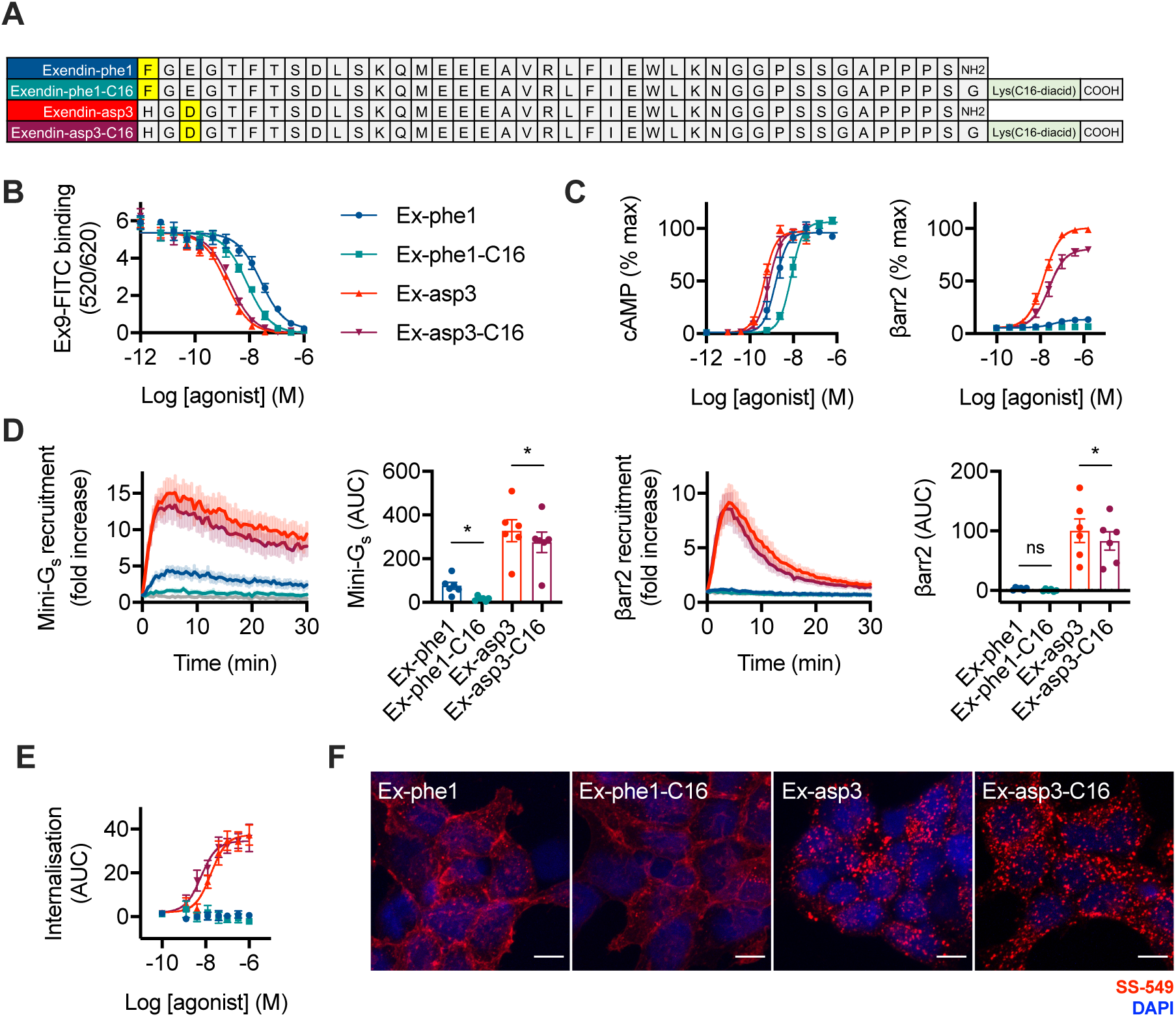
*In vitro* pharmacological properties of acylated biased GLP-1RAs. (**A**) Amino acid sequences in single letter code indicating position of C16 diacid conjugation. (**B**) Equilibrium binding of each ligand in competition with exendin(9-39)-FITC in HEK293-SNAP-GLP-1R cells, *n*=5. (**C**) Cyclic AMP (cAMP) production and β-arrestin-2 (βarr2) recruitment in CHO-K1-βarr2-EA-GLP-1R cells, 30-minute incubation, *n*=6, 4-parameter logistic fit of pooled data shown. (**D**) NanoBiT recruitment assays performed in HEK293T cells transiently transfected with GLP-1R-SmBiT and miniG_s_-LgBiT or LgBiT-β-arrestin-2, *n*=6, area-under-curve (AUC) compared using randomised block one-way ANOVA with Sidak’s test. (**E**) GLP-1R endocytosis measured in HEK293-SNAP-GLP-1R cells by DERET, indicated as AUC from kinetic traces shown in Supplementary Figure 1D, *n*=5, 4-parameter logistic fit of pooled data shown. (**F**) Representative images showing GLP-1R endocytosis in HEK293-SNAP-GLP-1R cells labelled with SNAP-Surface-549 (SS-549) prior to stimulation with indicated agonist (100 nM) for 30 minutes, *n*=3, scale bar: 8 µm. * p<0.05 by statistical test indicated in the text. Error bars indicate SEM.

Non-acylated exendin-phe1 is known to display markedly reduced recruitment of β-arrestin-1 and -2, whereas non-acylated exendin-asp3 is a full agonist in these pathways, with both ligands being full agonists for cAMP (12). To determine how acylation affects this established pattern of signal bias, we measured cAMP and β-arrestin-2 responses in PathHunter CHO-K1-βarr2-EA-GLP-1R cells. The expected signalling pattern was preserved (Figure 1C, Table 1), although both C16 ligands showed reduced efficacy for β-arrestin-2 recruitment compared to their non-acylated equivalents. Despite increased binding affinity, cAMP potency for exendin-phe1-C16 was reduced compared to that of exendin-phe1. Due to the undetectable β-arrestin-2 response with exendin-phe1-C16, quantification of signal bias was not possible either using the most commonly used approach (23, 24) or using an alternative method (22) (Supplementary Figure 1A).

**Table 1.**
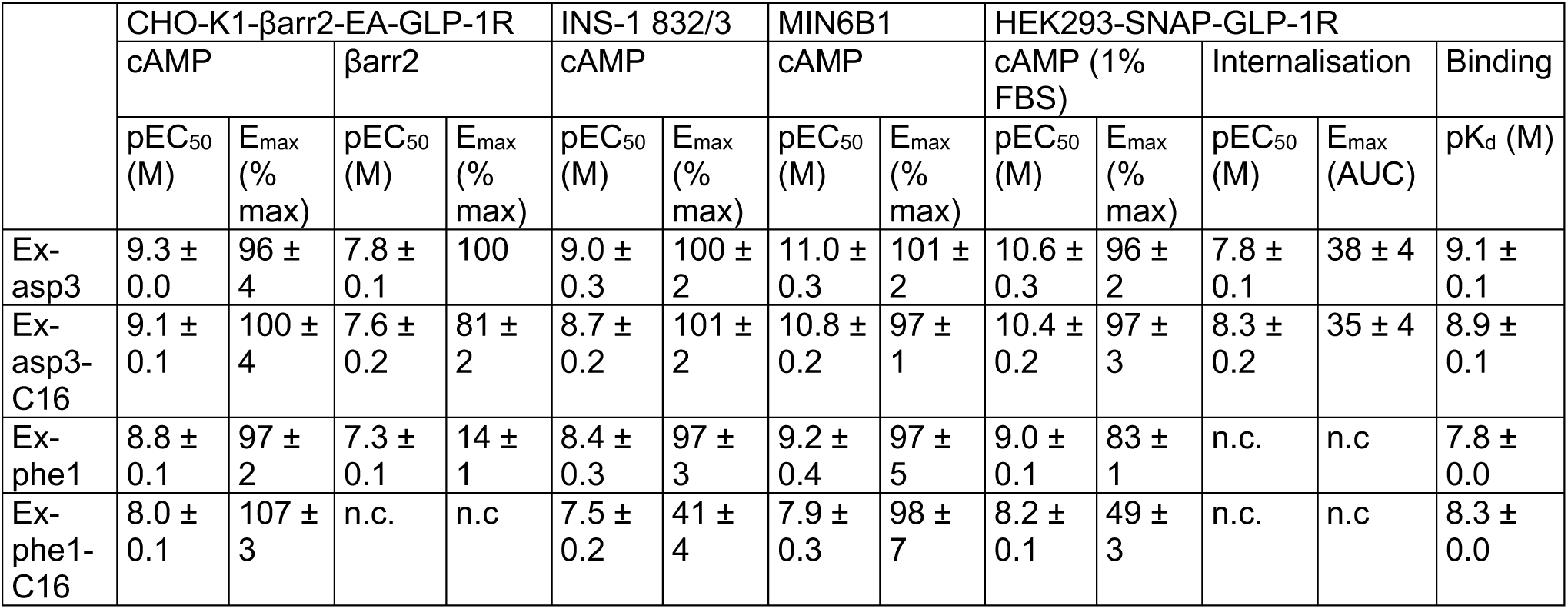
Binding, signalling and internalisation parameter estimates for ligands in this study. Signalling parameter measures were determined as follows: an initial 4-parameter fit was constructed for all full agonists, with globally constrained basal response, E_max_ and Hill slope, to establish the maximal response for the assay (note that for β-arrestin-2 measurement, Ex-asp3 was the only full agonist). Individual responses were normalised to the assay maximum and parameter estimates for each assay recalculated by 4-parameter fitting with globally constrained basal response and Hill slope, but no constraint to E_max_. For internalisation, curve fitting was performed similarly but without prior normalisation to a maximal response. Note that for β-arrestin-2, meaningful estimates for Ex-phe1-C16 could not be calculated (“n.c.”), as was also the case for both Ex-phe1 and Ex-phe1-C16 for internalisation. Average ± SEM values are reported.

| | CHO-K1-βarr2-EA-GLP-1R | | | | INS-1 832/3 | | MIN6B1 | | HEK293-SNAP-GLP-1R | | | | |
| --- | --- | --- | --- | --- | --- | --- | --- | --- | --- | --- | --- | --- | --- |
| | cAMP | | βarr2 | | cAMP | | cAMP | | cAMP (1% FBS) | | Internalisation | | Binding |
| | pEC<sub>50</sub> (M) | $E_{\max}$ (% max) | pEC<sub>50</sub> (M) | $E_{\max}$ (% max) | pEC<sub>50</sub> (M) | $E_{\max}$ (% max) | pEC<sub>50</sub> (M) | $E_{\max}$ (% max) | pEC<sub>50</sub> (M) | $E_{\max}$ (% max) | pEC<sub>50</sub> (M) | $E_{\max}$ (AUC) | pK<sub>d</sub> (M) |
| Ex-asp3 | 9.3 ± 0.0 | 96 ± 4 | 7.8 ± 0.1 | 100 | 9.0 ± 0.3 | 100 ± 2 | 11.0 ± 0.3 | 101 ± 2 | 10.6 ± 0.3 | 96 ± 2 | 7.8 ± 0.1 | 38 ± 4 | 9.1 ± 0.1 |
| Ex-asp3-C16 | 9.1 ± 0.1 | 100 ± 4 | 7.6 ± 0.2 | 81 ± 2 | 8.7 ± 0.2 | 101 ± 2 | 10.8 ± 0.2 | 97 ± 1 | 10.4 ± 0.2 | 97 ± 3 | 8.3 ± 0.2 | 35 ± 4 | 8.9 ± 0.1 |
| Ex-phe1 | 8.8 ± 0.1 | 97 ± 2 | 7.3 ± 0.1 | 14 ± 1 | 8.4 ± 0.3 | 97 ± 3 | 9.2 ± 0.4 | 97 ± 5 | 9.0 ± 0.1 | 83 ± 1 | n.c. | n.c. | 7.8 ± 0.0 |
| Ex-phe1-C16 | 8.0 ± 0.1 | 107 ± 3 | n.c. | n.c. | 7.5 ± 0.2 | 41 ± 4 | 7.9 ± 0.3 | 98 ± 7 | 8.2 ± 0.1 | 49 ± 3 | n.c. | n.c. | 8.3 ± 0.0 |

Signalling potencies for cAMP at rat and mouse GLP-1Rs was also tested in INS-1 832/3 and MIN6B1 insulinoma cells, respectively (Supplementary Figure 1B, C, Table 1). In these cells, which express the GLP-1R endogenously, the potency shift with exendin-phe1-C16 was more pronounced (up to 1000-fold reduction in MIN6B1 compared to exendin-asp3-C16), accompanied by reduced efficacy in INS-1 832/3 cells. These differences are more marked than for cells with GLP-1R exogenously expressed at high levels, e.g. the experiments shown in Figure 1C.

We also used NanoBiT complementation to monitor dynamic interactions of both β-arrestin-2 (25) and the mini-G_s_ protein probe (19) with the GLP-1R after stimulation with a maximal concentration of each ligand (Figure 1D). Both phe1 ligands showed markedly reduced efficacy for G_s_ recruitment when compared to asp3 ligands, but as this was accompanied by almost undetectable β-arrestin-2 recruitment, the phe1 ligands appear to be genuinely biased in favour of G protein signalling. In a separate set of experiments we compared mini-G_s_, -G_i_ and -G_q_ responses with each compound, confirming that G_s_ is the preferred G protein coupled to GLP-1R activation, but also revealing low magnitude G_q_ recruitment responses with both asp3 ligands (Supplementary Figure 1D,E).

Next, the four ligands were compared in HEK293-SNAP-GLP-1R cells for their propensity to induce GLP-1R internalisation, as measured by diffusion-enhanced resonance energy transfer (DERET) assay (26). Exendin-asp3 and exendin-asp3-C16 induced rapid internalisation, whereas none was detectable with either type of phe1 ligand (Figure 1E, Supplementary Figure 1F, Table 1). These results were corroborated by confocal microscopy (Figure 1F).

This initial *in vitro* characterisation demonstrates that the C16 C-terminal conjugation is well tolerated by exendin-asp3. Some differences were however observed with the pharmacology of exendin-phe1, which showed increased binding affinity but decreased signalling efficacy, thereby magnifying signalling differences between the two oppositely biased ligands.

### 3.2 Acylated biased GLP-1RAs show prolonged pharmacokinetics

Acylation prolongs peptide pharmacokinetics by promoting reversible binding to plasma proteins that are too large to undergo glomerular filtration, such as albumin. To determine the extent of binding of exendin-phe1-C16, exendin-asp3-C16 and the similarly designed exendin-4-C16 (Supplementary Figure 2A) to plasma proteins, we used the *ES*calate equilibrium shift assay (21). Results indicated that each acylated ligand exhibited a high and similar degree of binding to mouse and human plasma proteins (Table 2). In keeping with this, the presence of serum during the incubation did not differentially affect the signalling potency for each C16 ligand in GLP-1R cAMP assays compared to its non-acylated comparator (Supplementary Figure 2B); potency was in fact higher when serum was present for both acylated and non-acylated ligands, presumably due to reduced adsorption of peptide onto the microplate surface. In this system, an approximately 100-fold potency difference between exendin-phe1-C16 and exendin-asp3-C16 was observed, along with reduced efficacy for the former (Table 1).

**Table 2:** EScalate results demonstrating binding of each ligand to human and mouse plasma proteins. Data are expressed as mean ± SEM of percentage peptide unbound to plasma proteins, determined in duplicate.

| Compound | Unbound fraction ($f_u$) | |
| --- | --- | --- |
|  | Human plasma | Mouse plasma |
| Ex4-C16 | 2.3% ± 1.0% | 2.6% ± 0.1% |
| Ex-phe1-C16 | 2.1% ± 0.1% | 2.4% ± 0.3% |
| Ex-asp3-C16 | 2.2% ± 0.1% | 2.1% ± 0.3% |

In keeping with the anticipated pharmacokinetic effect resulting from binding to plasma proteins, after a single injection in mice, exendin-4-C16 remained detectable in the circulation for at least 72 hours, in contrast to non-acylated exendin-4 (Figure 2A). Similarly, exendin-phe1-C16 and exendin-asp3-C16 concentrations persisted for at least 72 hours (Figure 2B). We considered the possibility of whether GLP-1R-mediated clearance may play a role in pharmacokinetics once glomerular filtration is no longer a primary route of elimination, in a form of target-mediated drug disposal (TMDD) (27, 28), but the absence of large differences in circulating concentrations of exendin-phe1-C16 and exendin-asp3-C16 at 24 and 72 hours post-dosing argues against this possibility.

**Figure 2.**
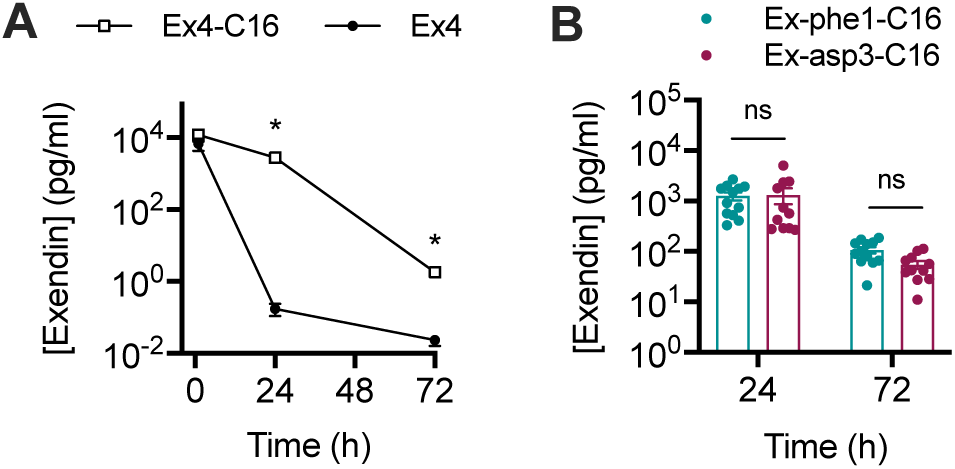
Pharmacokinetic properties of acylated biased GLP-1RAs. (**A**) Plasma concentration of exendin-4 (Ex4) or exendin-4-C16 (Ex4-C16) (both referred to as “Exendin” on the y-axis) in male C57Bl/6 mice after a single intraperitoneal injection of 100 nmol/kg agonist, *n*=8 per treatment, two-way repeat measures ANOVA on log-transformed data with Sidak’s test. (**B**) Plasma concentration of exendin-phe1-C16 or exendin-asp3-C16 in male C57Bl/6 mice after a single intraperitoneal injection of 100 nmol/kg agonist, *n*=12 (exendin-phe1-C16) or 11 (exendin-asp3-C16), two-way repeat measures ANOVA on log-transformed data with Sidak’s test. * p<0.05 by statistical test indicated in the text. Error bars indicate SEM.

Therefore, conjugation of a C16 diacid to the C-terminus of exendin-phe1 and exendin-asp3 results in a pair of pharmacokinetically advantageous and oppositely biased GLP-1RAs, allowing convenient assessment of the impact of signal bias and trafficking over multiple days.

### 3.3 Metabolic effects of biased GLP-1RAs are preserved after acylation

The primary therapeutic actions of GLP-1RAs are to improve glucose regulation and reduce appetite. We trialled a series of doses of exendin-phe1-C16 and exendin-asp3-C16 in lean C57Bl/6 mice to assess their *in vivo* performance over 72 hours. Both ligands exhibited effective glucoregulatory properties acutely (Supplementary Figure 3A), although exendin-phe1-C16 was less effective at 10 nmol/kg, which could reflect its reduced signalling potency. However, when reassessed at 72 hours, exendin-asp3-C16 no longer exhibited any detectable anti-hyperglycaemic efficacy, whereas both of the higher exendin-phe1-C16 doses remained effective. The acute anorectic effect of exendin-asp3-C16 was greater than that of exendin-phe1-C16, particularly at the 10 nmol/kg dose (Supplementary Figure 3B), although, interestingly, by 72 hours, the net calorie intake deficit with exendin-phe1-C16 at the highest dose was greater than that of exendin-asp3-C16. Dose response analysis of these data is displayed in Supplementary Figure 3C.

We also tested both compounds at an intermediate dose in obese mice fed a high fat diet for 3 months. Here, consistent patterns were observed, with equal glucose lowering seen 2 hours after dosing but a glycaemic advantage for exendin-phe1-C16 clearly demonstrated at 72 hours (Figure 3A). The anorectic differences were more subtle than in lean mice, with non-significant trends observed at 1 and 72 hours after dosing (Figure 3B). A non-significant trend was also seen for body weight loss, with exendin-phe1-C16-treated mice losing slightly more weight at 72 hours than those treated with exendin-asp3-C16 (Figure 3C).

**Figure 3.**
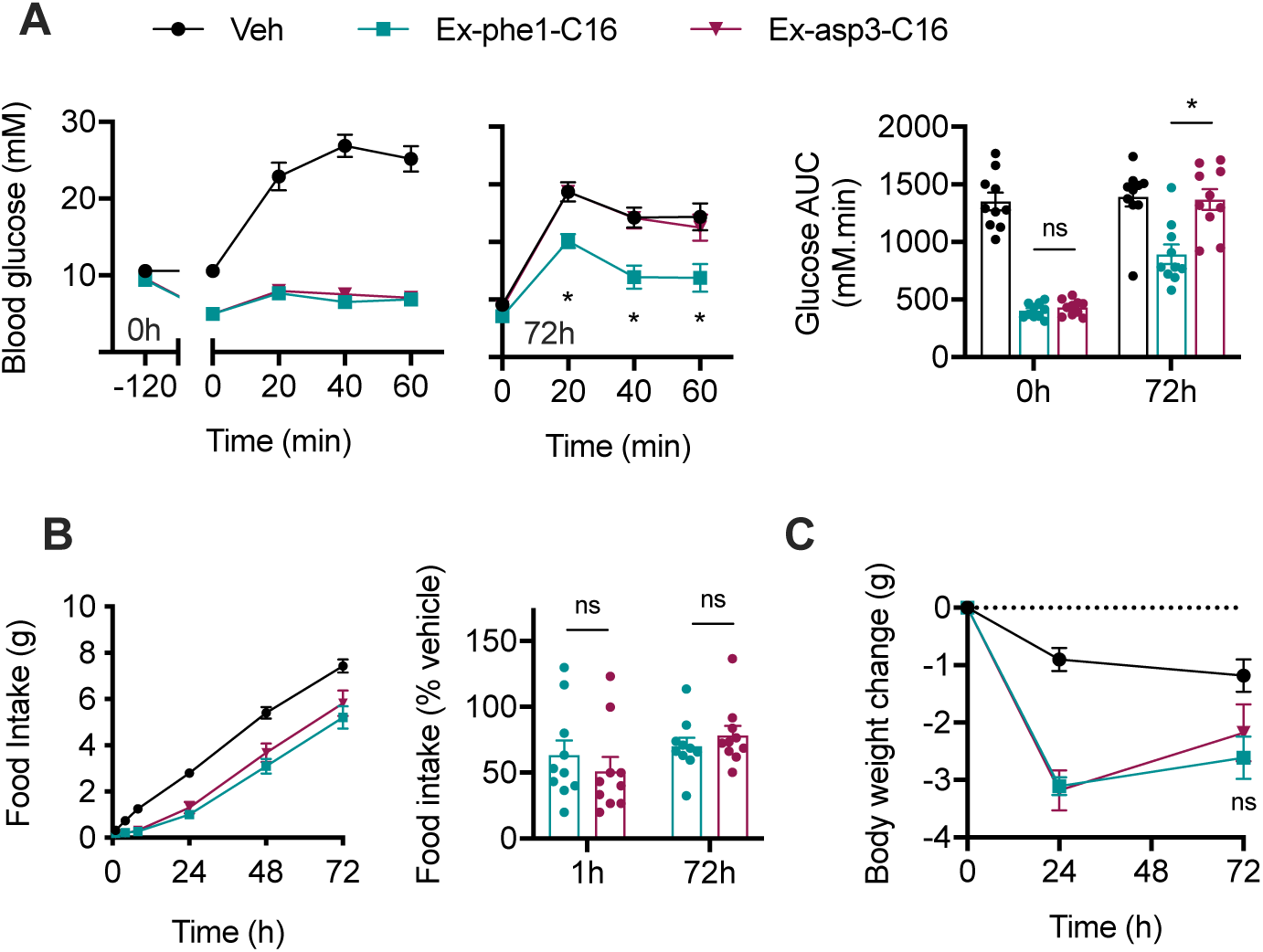
Sub-chronic effects of acylated biased GLP-1RAs in diet-induced obese mice. (**A**) Intra-peritoneal glucose tolerance tests (IPGTT, 2 g/kg glucose) performed 2 or 72 hours after IP administration of indicated -C16 agonist (10 nmol/kg) or vehicle (saline) in male DIO C57Bl/6 mice, *n*=10/group, time-points and AUCs both compared by two-way repeat measures ANOVA with Tukey’s test, with comparisons between exendin-phe1-C16 and exendin-asp3-C16 shown. (**B**) Cumulative food intake in male DIO C57Bl/6 mice after IP administration of indicated -C16 agonist (10 nmol/kg) or vehicle (saline), *n*=10/group, effect of each agonist to reduce food intake relative to vehicle at 1 and 72 hours is shown separately. (**C**) Body weight change in study shown in (B), two-way repeat measures ANOVA with Tukey’s test, comparisons between exendin-phe1-C16 and exendin-asp3-C16 are shown on the graph. * p<0.05 by statistical test indicated in the text. Error bars indicate SEM.

The apparent greater efficacy with exendin-phe1-C16 at 72 hours suggest that the impact of its biased pharmacology is preserved *in vivo*, despite its considerably lower net signalling potency. As observed previously, these effects primarily concerned its glucoregulatory properties.

### 3.4 Sustained administration of acylated biased GLP-1RAs

We performed a repeated administration study in high fat diet-induced obese mice to compare the therapeutic effects of exendin-phe1-C16 and exendin-asp3-C16 in a chronic setting. Mice were injected every 72 hours over 15 days; the dose was doubled after the first 3 injections to counteract adaptive mechanisms typically seen in rodents treated with GLP-1RAs which limit weight loss (30–32). Over the course of the study, the trends observed in the single dose administration studies became more apparent, with a progressively greater anorectic effect observed with exendin-phe1-C16, along with corresponding divergence in body weight (Figure 4A, B). Glucose tolerance assessed 72 hours after the final dose confirmed the expected advantage of exendin-phe1-C16 over exendin-asp3-C16 (Figure 4C).

**Figure 4.**
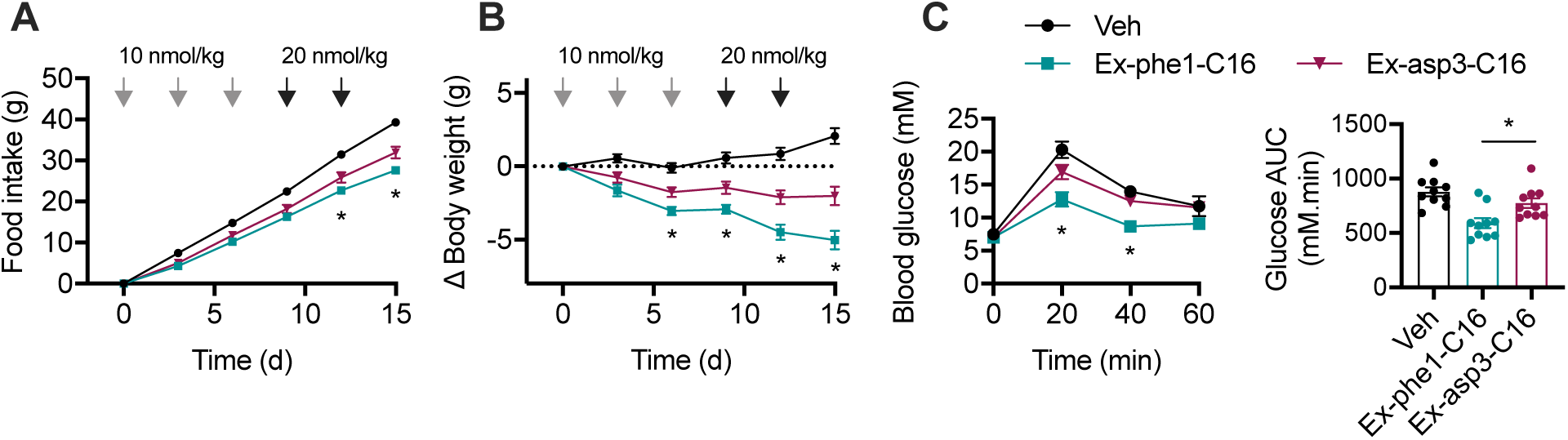
Effects of chronic administration of biased GLP-1RAs in diet-induced obese mice. (**A**) Cumulative food intake in male DIO C57Bl/6 mice after IP administration of -C16 agonist every 72 hours or vehicle (saline), *n*=10/group, two-way repeat measures ANOVA with Tukey’s test showing comparison between exendin-phe1-C16 and exendin-asp3-C16. (**B**) Change in body weight during study shown in (A), two-way repeat measures ANOVA with Tukey’s test showing comparison between exendin-phe1-C16 and exendin-asp3-C16. (**C**) IPGTT (2 g/kg glucose) performed 72 hours after the final agonist dose, time-points compared by two-way repeat measures ANOVA with Tukey’s test (comparison between exendin-phe1-C16 and exendin-asp3-C16 is shown) and AUC compared by one-way ANOVA with Tukey’s test (comparison between exendin-phe1-C16 and exendin-asp3-C16 is shown). * p<0.05 by statistical test indicated in the text. Error bars indicate SEM.

Thus, the metabolic benefits of the biased GLP-1RA exendin-phe1-C16 were preserved on repeated administration, and became progressively more apparent over time.

## 4 Discussion

In the present study, we have developed and tested two oppositely biased GLP-1RAs bearing C-terminal C16 diacid chains, which confer extended pharmacokinetic profiles. The major observation from our data is that, despite at least large reduction in cAMP signalling potency (10- to 1000-fold, depending on cell system used), exendin-phe1-C16 is markedly more efficacious *in vivo* than oppositely biased exendin-asp3-C16. This finding builds on earlier reports highlighting the advantageous glucoregulatory properties of lower affinity biased GLP-1RAs which favour cAMP signalling over β-arrestin recruitment (11, 12), but with a substantially greater disconnect between acute *in vitro* potency and anti-hyperglycaemic properties than for earlier compounds.

Some interesting *in vitro* observations arise from this work, particularly regarding comparisons of acylated *versus* non-acylated ligands. Firstly, we found that the binding affinity of exendin-phe1-C16 was approximately 3 times greater than that of non-acylated exendin-phe1, yet its cAMP signalling potency was between 5 and 20 times (for CHO-K1 and MIN6B1 cells, respectively) lower. One possible explanation for this would be that the acyl chain disrupts normal ligand-receptor interactions in a manner which permits ligand binding but orientates the ligand so that its N-terminus fails to engage optimally with the receptor core, leading to reduced levels of activation. Structural and computational studies will likely be required to reveal if is this is the case, and may be hampered by the lack of confidence about the position of the exendin-4 C-terminus when bound to GLP-1R (33). An alternative possibility, tentatively supported by recent reports that active GLP-1Rs segregate into cholesterol-rich nanodomains within the plasma membrane (17, 29), is that the fatty acid preferentially directs the ligand to bind to subpopulations of GLP-1Rs residing in membrane regions with reduced enrichment of signalling effectors on the cytoplasmic side.

Addition of a C16 diacid moiety, as expected from other ligands tested as part of the preclinical development of C18 diacid-containing semaglutide (34, 35), provided a high degree of binding to both mouse and human plasma proteins. The slightly lower affinity of the investigated compounds (2.08% – 2.33% in human plasma) compared to Liraglutide, containing a C16 fatty acid moiety (0.51% in human plasma (21)), may be attributable to the shorter linker between the peptide and the albumin binding group. In combination with resistance of exendin peptides to degradation by a variety of enzymes including dipeptidyl dipeptidase-4 and neutral endopeptidases (36), the two primary elimination routes for GLP-1 peptides are avoided, providing a basis for substantially extended pharmacokinetics in mice, with even longer protraction expected to be obtained in humans.

Our evaluation of the pharmacodynamic performance of biased GLP-1RAs found clear evidence of improved anti-hyperglycaemic efficacy for exendin-phe1-C16. This is in spite of substantially reduced acute signalling potency of this molecule, highlighting how standard *in vitro* approaches may fail to identify optimal agonist characteristics. Even though coupling to G_s_ recruitment was markedly reduced, it can be speculated that the virtual absence of β-arrestin recruitment, and of GLP-1R internalisation, allow prolongation of signalling despite an initial deficit, as previously described (12). Interestingly, recent work has suggested that β-arrestins play a minimal role in controlling GLP-1R endocytosis (17), and may not diminish acute insulin secretory responses from pancreatic beta cells (9, 37). However, the latter observation needs to be evaluated under conditions of sustained exposure to pharmacokinetically optimised GLP-1RAs, in line with the appropriate timescales for glucoregulatory benefits of biased GLP-1RAs, which tend to emerge after a number of hours. Interestingly, GLP-1R internalisation was reported to be G_q_-dependent (38), and with this in mind, it may be relevant that we detected G_q_ recruitment to the GLP-1R after treatment with fast-internalising -asp3 ligands.

It is notable that the differences in physiological response entrained by oppositely biased ligands concerned primarily their glucoregulatory effects, with smaller differences observed on feeding behaviour. A similar pattern has previously been noted with other biased GLP-1RAs (11, 12), and this divergence has not yet been satisfactorily explained. Differential access to anorectic neurons within the central nervous system due to altered GLP-1R-mediated carriage across the blood brain barrier (39), as well as differential actions of biased ligands on different cell types [“tissue bias” (40)] remaining two realistic possibilities to be explored. Nevertheless, it should be highlighted that, in the present report, when administered chronically, exendin-phe1-C16 resulted in superior cumulative reductions in food intake and greater weight loss than exendin-asp3-C16. As exendin-phe-C16 tended to exert a milder appetite suppressive effect in the acute setting, enhanced tolerability might result whilst still achieving better weight loss and anti-hyperglycaemic efficacy.

In this study we did not compare these ligands against class-leading GLP-1RAs such as semaglutide and dulaglutide, and therefore their true therapeutic potential remains unclear. Nevertheless, our results clearly show that agonists displaying weak acute signalling efficacy and potency can be surprisingly effective for therapeutically important readouts. Understanding the molecular events which underpin their unusual pharmacology may also shed light on new ways to control GLP-1R behaviours in the future.

## Abbreviations

AUC: Area under curve
BSA: Bovine serum albumin
cAMP: Cyclic adenosine monophosphate
FITC: Fluorescein isothiocyanate
GLP-1RA: Glucagon-like peptide-1 receptor agonist
HBSS: Hank’s buffered salt solution
HTRF: Homogenous time-resolved fluroescence
LC/MSMS: Liquid chromatography / tandem mass spectrometry
n.c.: Not calculable
PBS: Phosphate-buffered saline
SEM: Standard error of the mean
T2D: Type 2 diabetes
TR-FRET: Time-resolved Förster resonance energy transfer

## Acknowledgements

We are grateful to Prof Nevin Lambert (Medical College of Georgia) for providing miniG-LgBiT plasmids. This work was funded by an MRC project grant to B.J., A.T., S.R.B. and G.A.R. The Section of Endocrinology and Investigative Medicine is funded by grants from the MRC, BBSRC, NIHR, an Integrative Mammalian Biology (IMB) Capacity Building Award, an FP7-HEALTH-2009-241592 EuroCHIP grant and is supported by the NIHR Biomedical Research Centre Funding Scheme. The views expressed are those of the author(s) and not necessarily those of the funder. B.J. was also supported for this work by the Academy of Medical Sciences, Society for Endocrinology and an EPSRC capital award. G.A.R. was supported by Wellcome Trust Investigator (212625/Z/18/Z) Awards, MRC Programme (MR/R022259/1) and Experimental Challenge Grant (DIVA, MR/L02036X/1), and Diabetes UK (BDA/11/0004210, BDA/15/0005275, BDA 16/0005485) grants. This work has received support from the EU/EFPIA/Innovative Medicines Initiative 2 Joint Undertaking (RHAPSODY grant No 115881) to G.A.R.

---

**Supplementary Figure 1.**
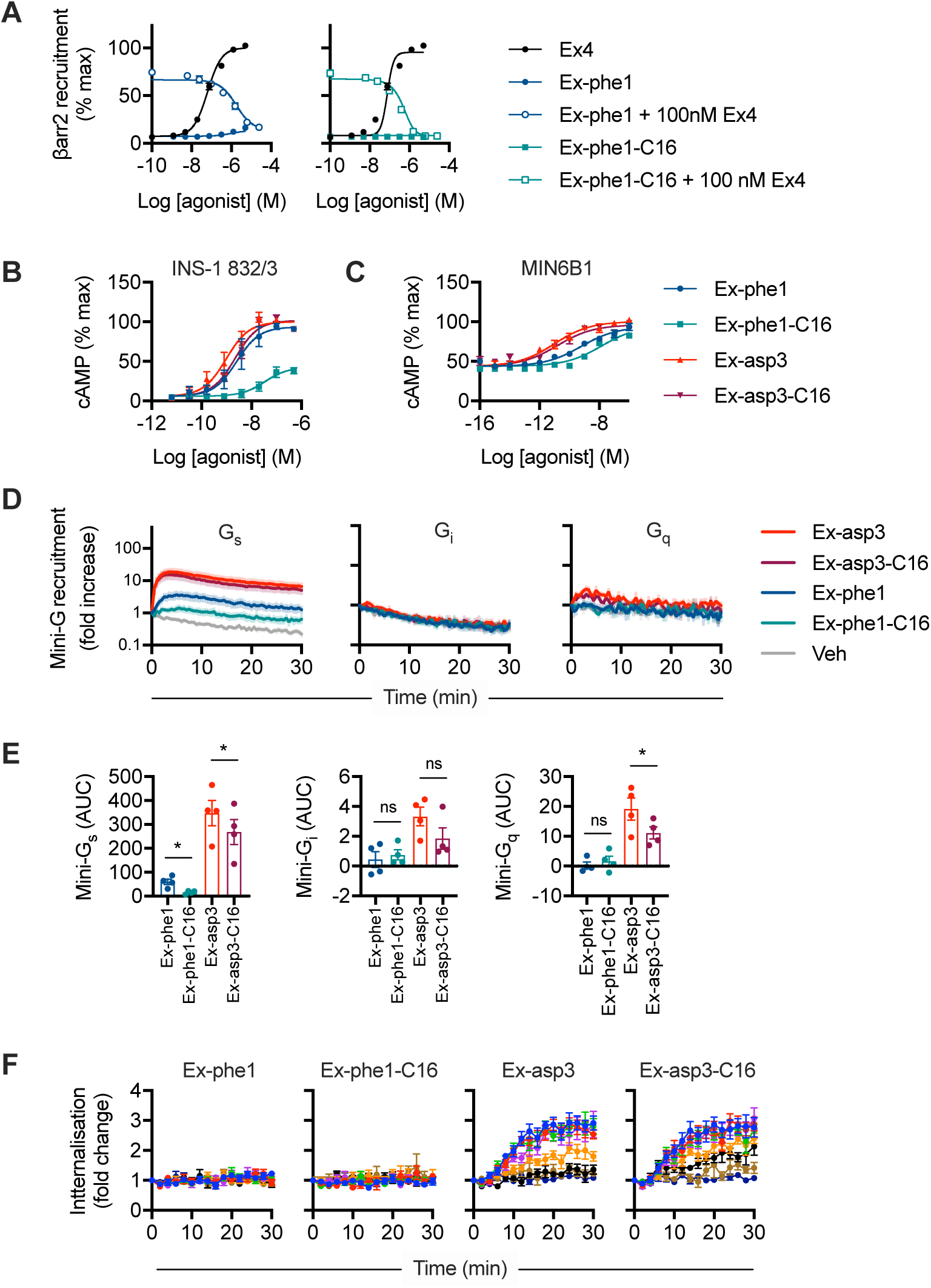
Additional *in vitro* data. (**A**) β-arrestin-2 recruitment in CHO-K1-GLP-1R β-arrestin-2-EA cells treated with indicated ligand or ligand combination, 30-minute incubation, *n*=4, curve fitting of pooled data using method of Stahl *et al*. (22). (**B**) cAMP responses in INS-1 832/3 cells treated with indicated agonist in presence of 500 µM IBMX, 10-minute incubation, *n*=4, 4-parameter logistic fit of pooled data shown. (**C**) cAMP responses in MIN6B1 cells treated with indicated agonist in presence of 500 µM IBMX, 5-minute incubation, *n*=5, 4-parameter logistic fit of pooled data shown. (**D**) NanoBiT recruitment assays performed in parallel in HEK293T cells transiently transfected with GLP-1R-smBiT and miniG_s_-lgBiT, miniG_i_-lgBiT or miniG_q_-lgBiT, stimulated with vehicle of indicated agonist (1 µM), note logarithmic scale to allow comparison of each miniG protein response, *n*=4. (**E**) AUCs from (D), calculated after subtracting vehicle response on linear scale, compared using randomised block one-way ANOVA with Sidak’s test. (**F**) DERET traces indicating GLP-1R internalisation in HEK293-SNAP-GLP-1R cells, *n*=5, pertains to Figure 1E. * p<0.05 by statistical test indicated in the text. Error bars indicate SEM.

**Supplementary Figure 2.**
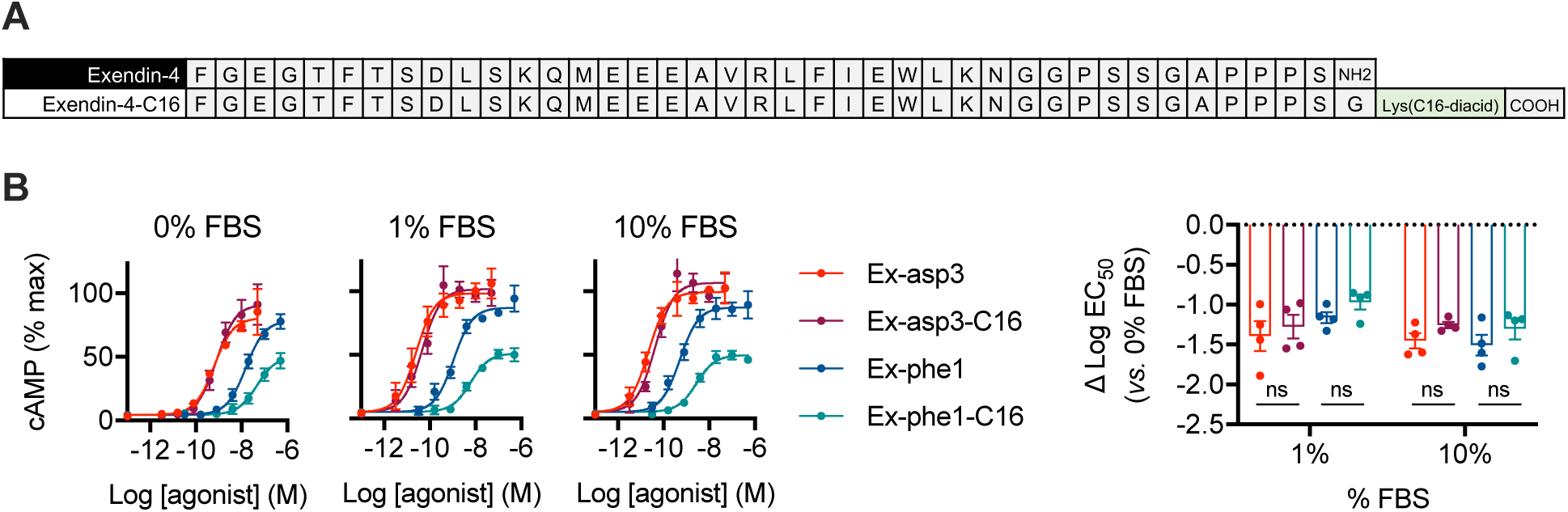
Additional data concerning protein binding of -C16 ligands. (**A**) Amino acid sequences of exendin-4 and exendin-4-C16 in single letter code. (**B**) cAMP responses in HEK293-SNAP-GLP-1R cells in presence of indicated concentration of FBS, 30-minute incubation, *n*=4, 4-parameter logistic fit of pooled data shown, and effect of ± FBS is determined by subtracting logEC_50_ values for each ligand; all comparisons between acylated and non-acylated ligands are non-significant by two-way repeat measures ANOVA with Tukey’s test. Error bars indicate SEM.

**Supplementary Figure 3.**
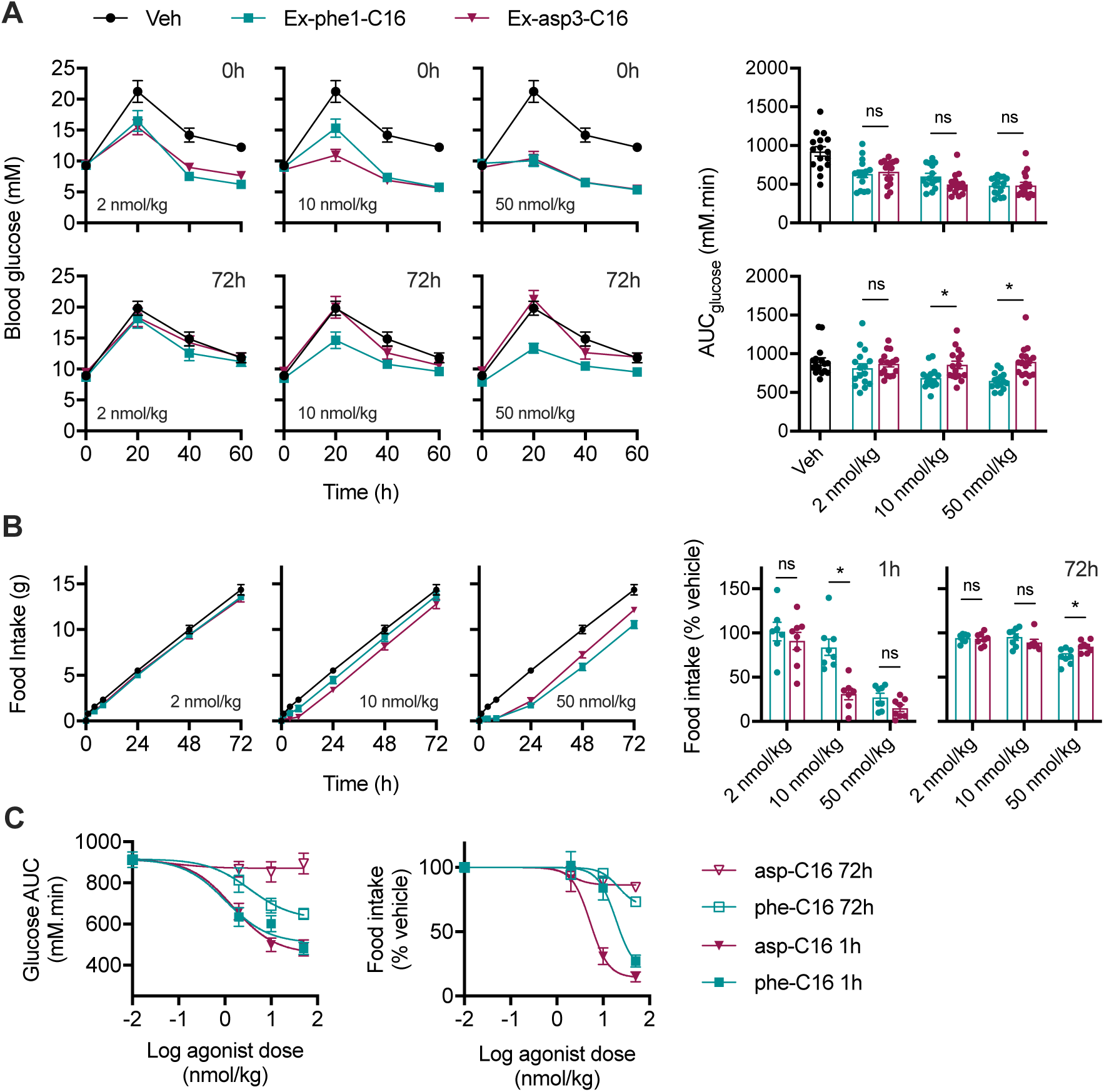
Dose range studies in lean mice. (**A**) IPGTTs (2 g/kg glucose) performed concurrently with or 72 hours after IP administration of indicated dose of -C16 agonist in lean, male C57Bl/6 mice, *n*=15-16/group, AUCs compared by two-way ANOVA with Sidak’s test. (**B**) Cumulative food intake over 72 hours after IP administration of indicated dose of -C16 agonist in lean, male C57Bl/6 mice, *n*=7-8/group, 1- and 72-hour data plotted separately, two-way repeat measures ANOVA with Tukey’s test. (**C**) Data from (A) and (B) displayed as a dose response analysis with 4-parameter logistic fit shown. * p<0.05 by statistical test indicated in the text. Error bars indicate SEM.

## Supplementary data EScalate plasma protein binding assay

Plasma protein binding of Ex4-C16, Ex-asp3-C16 and Ex-phe1-C16 was determined using the 3BP *EScalate* Equilibrium Shift Assay. In short, the shift of the binding equilibrium of the compound to HSA-coated beads following addition of plasma at various dilutions was analysed. From this concentration-dependent shift, the apparent dissociation constants for binding to HSA on the beads and binding to plasma proteins can be calculated. From the apparent dissociation constant to plasma proteins, the fraction that is not bound to plasma proteins (fraction unbound, f_u_) can be calculated.

### Materials

Acetonitrile super gradient grade was obtained from VWR, Darmstadt, Germany. Trifluoroacetic acid LC-MS grade was delivered by Thermo Fisher Scientific, Schwerte, Germany.

Formic acid LC-MS grade and Dimethyl sulfoxide (DMSO) puriss. grade was purchased from Sigma Aldrich, Taufkirchen, Germany.

Water was purified using a Millipore milli-Q A-10 water purification system. Dulbecco’s phosphate buffered saline (PBS) without Mg^2+^ and Ca^2+^ was obtained from Biochrom GmbH, Berlin, Germany.

### Stock solutions

1 mM stock solution Ex4-C16: 1.65 mg of the compound was dissolved in 356 µL DMSO. 1 mM stock solution Ex4-asp3-C16: 1.85 mg of the compound was dissolved in 400 µL DMSO.

1 mM stock solution Ex4-phe1-C16: 1.61 mg of the compound was dissolved in 346 µL DMSO.

24 µM stock solutions: 24 µL of the 1 mM stock solution was diluted with 976 µL PBS.

### Plasma stocks

#### Human plasma

Pooled normal K2EDTA plasma, BioIVT, Burges Hill, Great Britain

#### Mouse plasma CD1(ICR)

Pooled CD-1 mouse K2EDTA plasma, BioIVT, Burges Hill, Great Britain

### Plasma dilutions

Following plasma dilutions were prepared for each assay.

**Table 1.** Preparation of plasma dilutions

| Plasma dilution | Plasma volume | PBS volume |
| --- | --- | --- |
| 50% | 200 μl | 200 μl |
| 16.7% | 66.7 μl | 333 μl |
| 5.6% | 22.2 μl | 378 μl |
| 1.9% | 7.41 μl | 393 μl |
| 0.62% | 2.47 μl | 398 μl |

### *EScalate* Equilibrium Shift Assay

Lot: 538-158

The EScalate assay consists of five well lanes with 6 wells containing following concentration of immobilized HSA: 0 µM, 11.4 µM, 20.6 µM, 37.0 µM, 66.7 µM, 120 µM in 85 µl PBS.

### Sample generation

The Equilibrium Shift Assay samples were generated by addition of 30 µl of the prepared plasma dilutions (50%, 16.7%, 5.6%, 1.9% and 0.62%) and 5 µl of the 24 µM compound stock solution to each well.

A binding protein concentration of 600 µM was assumed for all plasma samples.

After 1 hour of incubation at room temperature under gentle shaking, the kit plate was centrifuged to separate the HSA immobilized on beads from the free plasma proteins. 50 µL of the supernatant were taken as sample from each vial.

For calibration purposes, additional compound solutions in PBS were prepared by diluting the 24 µM stock solution as following:

**Table 2.**
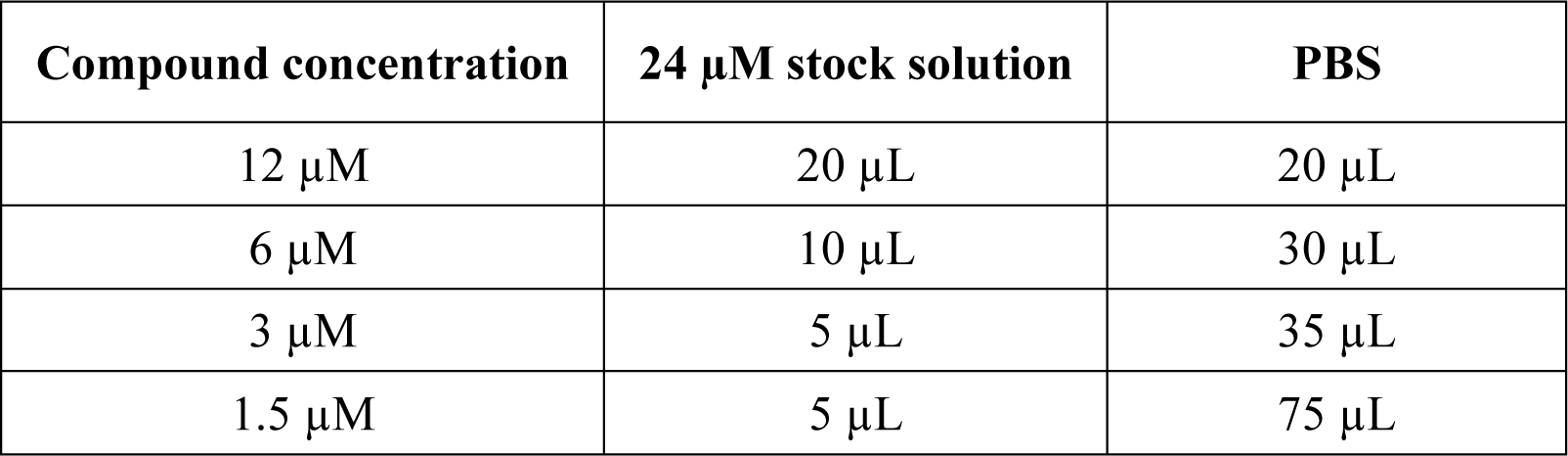
Preparation of calibration spiking solutions

Calibration samples were prepared by spiking 120 µL of the lowest plasma / protein dilution with 5 µL of the compound spiking solution. The calibration samples were incubated together with the assay samples.

### Sample preparation

Each sample was mixed with 2 µL of the 24 µM stock solution of the chosen internal standard and 250 µL of acidified acetonitrile (containing 1% trifluoroacetic acid) to precipitate the plasma proteins. The samples were centrifuged and 100 µL of the supernatants were diluted with 100 µL of 0.1% aqueous formic acid.

**Table 3.**
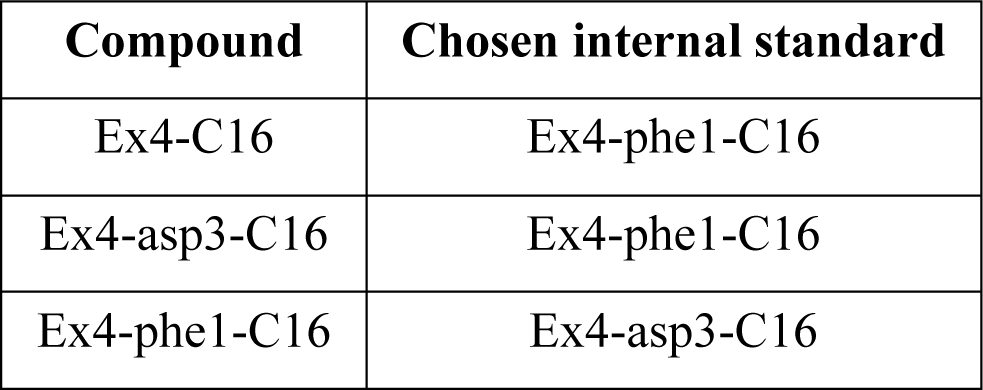
Assignment of internal standards to the test items

| **Compound** | **Chosen internal standard** |
| --- | --- |
| Ex4-C16 | Ex4-phe1-C16 |
| Ex4-asp3-C16 | Ex4-phe1-C16 |
| Ex4-phe1-C16 | Ex4-asp3-C16 |

### LC-MS analysis

The samples were analyzed using an Agilent 1290 UHPLC system coupled to an Agilent 6470 triple quadrupole mass spectrometer.

### Chromatographic parameters

| Compound name | Retention time | Fragmentor | Transition / Collision Energy |
| --- | --- | --- | --- |
| Ex4-C16 | 1.75 min | 190 V | Quantifier: 929.3 → 851.8 @ 22V<br>Qualifier: 929.3 → 129.2 @ 43 V |
| Ex4-asp3-C16 | 1.78 min | 160 V | Quantifier: 926.5 → 946.8 @ 31 V<br>Qualifier: 926.5 → 851.8 @ 21 V |
| Ex4-phe1-C16 | 1.97 min | 190 V | Quantifier: 1164.0 → 851.8 @ 20 V<br>Qualifier: 1164.0 → 754.3 @ 50 V |

### 1.1 Data analysis

A quadratic calibration was performed from 0.015 µM to 1 µM using the prepared calibration samples followed by quantitation of the compounds in all samples using the lowest plasma dilution as matrix.

From the determined compound concentrations (APA) of all sample data, the fitting procedure for both dissociation constants was performed according to the following equation. Function:

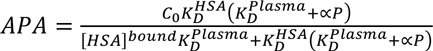

where c_0_ is the total concentration of drug in the incubation, K_D_^HSA^ is the dissociation constant of the compound from immobilized HSA on beads, K_D_^Plasma^ is the dissociation constant of the compound from plasma proteins in solution, [HSA]^bound^ is the concentration in solution of HSA bound to the beads, P is the concentration of the binding protein in plasma and α is the applied dilution factor.

Then, the free fraction of the compound in plasma was calculated as follows (assuming a binding protein concentration of 600 µM for all plasma samples).

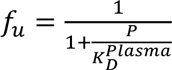

To evaluate the assay a Total Quality Index (TQI) was calculated to evaluate the consistency with the applied binding model, uncertainty of the analysis method and differences in compound recovery at different plasma dilutions. The index can take values between zero and ten. A value of 8 or higher is indicating good assay performance.

### Results

The results are given in Table 4.

**Table 4.** Unbound fraction in plasma of different species, determined apparent K_D_ values for the binding to plasma proteins and calculated Total Quality Indices

| Compound | Species | Unbound fraction | App. $K_D$ [μM] | TQI |
| --- | --- | --- | --- | --- |
| Ex4-C16 | Human | 2.16% / 2.49% | 13.4 / 15.5 | 9.3 / 9.5 |
| Ex4-C16 | Mouse | 2.59% / 2.57% | 15.8 / 15.8 | 9.3 / 9.3 |
| Ex4-asp3-C16 | Human | 2.06% / 2.10% | 12.6 / 12.8 | 9.3 / 9.5 |
| Ex4-asp3-C16 | Mouse | 2.32% / 2.43% | 14.2 / 14.8 | 9.3 / 8.9 |
| Ex4-phe1-C16 | Human | 2.16% / 2.18% | 13.4 / 13.4 | 9.3 / 9.5 |
| Ex4-phe1-C16 | Mouse | 2.13% / 2.08% | 12.3 / 12.8 | 8.5 / 9.1 |

The Total Quality Index (TQI) is an artificial value to describe the assay performance. It ranges from 1 to 10 and covers the consistency with the applied binding model, uncertainty of the analysis method and differences in compound recovery at different plasma dilutions. The index can take values between zero and ten. It is calculated as the mean of different quality indices which can also take values between zero and ten (see Table 5). A value of 8 or higher is indicating good assay performance.

**Table 5.** Description of used quality indices

| Quality index | Description |
| --- | --- |
| Recovery | Significant differences in the recovery rate of the different plasma dilution samples may lead to a distortion of the assay results. Homogenous recovery lead to high QI values. |
| Average QI | Each plasma dilution sample series shall fit to the theoretical model of exclusive single non-covalent binding of the compound to HSA. Poor fit will lead to low QI values. The mean of the quality indices of all plasma dilutions thus describes the applicability of the binding model. |
| 95% confidence interval of $f_u$ | Large intra-assay confidence interval shows poor assay performance |
| Number of outlier | High number of statistical outlier show poor assay performance |
| $f_b/f_u$ distance | The mean deviation between measured and calculated $f_b/f_u$ values for each plasma dilution series. Higher deviations indicate poor model fit. |

## References

1. Andersen A, Lund A, Knop FK, Vilsbøll T. Glucagon-like peptide 1 in health and disease. Nat Rev Endocrinol. 2018 Jul;14(7):390–403.

2. Mann JFE, Ørsted DD, Brown-Frandsen K, Marso SP, Poulter NR, Rasmussen S, et al. Liraglutide and Renal Outcomes in Type 2 Diabetes. N Engl J Med. 2017 Aug 31;377(9):839–48.

3. Bethel MA, Patel RA, Merrill P, Lokhnygina Y, Buse JB, Mentz RJ, et al. Cardiovascular outcomes with glucagon-like peptide-1 receptor agonists in patients with type 2 diabetes: a meta-analysis. Lancet Diabetes Endocrinol. 2018 Feb;6(2):105–13.

4. Zheng SL, Roddick AJ, Aghar-Jaffar R, Shun-Shin MJ, Francis D, Oliver N, et al. Association Between Use of Sodium-Glucose Cotransporter 2 Inhibitors, Glucagon-like Peptide 1 Agonists, and Dipeptidyl Peptidase 4 Inhibitors With All-Cause Mortality in Patients With Type 2 Diabetes: A Systematic Review and Meta-analysis. JAMA. American Medical Association; 2018 Apr 17;319(15):1580–91.

5. Drucker DJ, Philippe J, Mojsov S, Chick WL, Habener JF. Glucagon-like peptide I stimulates insulin gene expression and increases cyclic AMP levels in a rat islet cell line. Proc Natl Acad Sci USA. 1987 May;84(10):3434–8.

6. Hayes MR, Leichner TM, Zhao S, Lee GS, Chowansky A, Zimmer D, et al. Intracellular signals mediating the food intake-suppressive effects of hindbrain glucagon-like peptide-1 receptor activation. Cell Metab. 2011 Mar 2;13(3):320–30.

7. Buenaventura T, Kanda N, Douzenis PC, Jones B, Bloom SR, Chabosseau P, et al. A Targeted RNAi Screen Identifies Endocytic Trafficking Factors That Control GLP-1 Receptor Signaling in Pancreatic β-Cells. Diabetes. 2018 Mar;67(3):385–99.

8. Fletcher MM, Halls ML, Zhao P, Clydesdale L, Christopoulos A, Sexton PM, et al. Glucagon-like peptide-1 receptor internalisation controls spatiotemporal signalling mediated by biased agonists. Biochem Pharmacol. 2018 Oct;156:406–19.

9. Sonoda N, Imamura T, Yoshizaki T, Babendure JL, Lu J-C, Olefsky JM. Beta-Arrestin-1 mediates glucagon-like peptide-1 signaling to insulin secretion in cultured pancreatic beta cells. Proc Natl Acad Sci USA. 2008 May 6;105(18):6614–9.

10. Koole C, Wootten D, Simms J, Miller LJ, Christopoulos A, Sexton PM. Differential Impact of Amino Acid Substitutions on Critical Residues of the Human Glucagon-Like Peptide-1 Receptor (GLP-1R) Involved in Peptide Activity and Small Molecule Allostery. J Pharmacol Exp Ther. 2015 Jan 28.

11. Zhang H, Sturchler E, Zhu J, Nieto A, Cistrone PA, Xie J, et al. Autocrine selection of a GLP-1R G-protein biased agonist with potent antidiabetic effects. Nat Commun. 2015;6:8918.

12. Jones B, Buenaventura T, Kanda N, Chabosseau P, Owen BM, Scott R, et al. Targeting GLP-1 receptor trafficking to improve agonist efficacy. Nat Commun. Nature Publishing Group; 2018 Apr 23;9(1):1602.

13. Hager MV, Johnson LM, Wootten D, Sexton PM, Gellman SH. β-Arrestin-Biased Agonists of the GLP-1 Receptor from β-Amino Acid Residue Incorporation into GLP-1 Analogues. J Am Chem Soc. 2016 Nov 4.

14. Chen X, Mietlicki-Baase EG, Barrett TM, McGrath LE, Koch-Laskowski K, Ferrie JJ, et al. Thioamide Substitution Selectively Modulates Proteolysis and Receptor Activity of Therapeutic Peptide Hormones. J Am Chem Soc. 2017 Nov 13.

15. Fremaux J, Venin C, Mauran L, Zimmer R, Koensgen F, Rognan D, et al. Ureidopeptide GLP-1 analogues with prolonged activity in vivo via signal bias and altered receptor trafficking. Chem Sci. Royal Society of Chemistry; 2019.

16. Simonsen L, Holst JJ, Deacon CF. Exendin-4, but not glucagon-like peptide-1, is cleared exclusively by glomerular filtration in anaesthetised pigs. Diabetologia. 2006 Apr;49(4):706–12.

17. Buenaventura T, Bitsi S, Laughlin WE, Burgoyne T, Lyu Z, Oqua AI, et al. Agonist-induced membrane nanodomain clustering drives GLP-1 receptor responses in pancreatic beta cells. Titchenell P, editor. PLoS Biol. 2019 Aug;17(8):e3000097.

18. Widmann C, Dolci W, Thorens B. Agonist-induced internalization and recycling of the glucagon-like peptide-1 receptor in transfected fibroblasts and in insulinomas. Biochem J. 1995 Aug 15;310 (Pt 1):203–14.

19. Wan Q, Okashah N, Inoue A, Nehmé R, Carpenter B, Tate CG, et al. Mini G protein probes for active G protein-coupled receptors (GPCRs) in live cells. J Biol Chem. American Society for Biochemistry and Molecular Biology; 2018 Mar 9;:jbc.RA118.001975.

20. Kroeze WK, Sassano MF, Huang X-P, Lansu K, McCorvy JD, Giguère PM, et al. PRESTO-Tango as an open-source resource for interrogation of the druggable human GPCRome. Nat Struct Mol Biol. Nature Publishing Group; 2015 May;22(5):362–9.

21. Ungewiss J, Gericke S, Boriss H. Determination of the Plasma Protein Binding of Liraglutide Using the EScalate* Equilibrium Shift Assay. J Pharm Sci. 2019 Mar;108(3):1309–14.

22. Stahl EL, Zhou L, Ehlert FJ, Bohn LM. A novel method for analyzing extremely biased agonism at G protein-coupled receptors. Mol Pharmacol. 2015 May;87(5):866–77.

23. Kenakin T, Watson C, Muniz-Medina V, Christopoulos A, Novick S. A simple method for quantifying functional selectivity and agonist bias. ACS Chem Neurosci. 2012 Mar 21;3(3):193–203.

24. van der Westhuizen ET, Breton B, Christopoulos A, Bouvier M. Quantification of ligand bias for clinically relevant β2-adrenergic receptor ligands: implications for drug taxonomy. Mol Pharmacol. 2014 Mar;85(3):492–509.

25. Dixon AS, Schwinn MK, Hall MP, Zimmerman K, Otto P, Lubben TH, et al. NanoLuc Complementation Reporter Optimized for Accurate Measurement of Protein Interactions in Cells. ACS Chem Biol. 2016 Feb 19;11(2):400–8.

26. Levoye A, Zwier JM, Jaracz-Ros A, Klipfel L, Cottet M, Maurel D, et al. A Broad G Protein-Coupled Receptor Internalization Assay that Combines SNAP-Tag Labeling, Diffusion-Enhanced Resonance Energy Transfer, and a Highly Emissive Terbium Cryptate. Front Endocrinol (Lausanne). 2015;6:167.

27. Gibiansky L, Gibiansky E. Target-mediated drug disposition model: approximations, identifiability of model parameters and applications to the population pharmacokinetic-pharmacodynamic modeling of biologics. Expert Opin Drug Metab Toxicol. 2009 Jul;5(7):803–12.

28. Gao W, Jusko WJ. Target-mediated pharmacokinetic and pharmacodynamic model of exendin-4 in rats, monkeys, and humans. Drug Metab Dispos. 2012 May;40(5):990–7.

29. Ast J, Arvaniti A, Fine NHF, Nasteska D, Ashford FB, Stamataki Z, et al. LUXendins reveal endogenous glucagon-like peptide-1 receptor distribution and dynamics. bioRxiv. Cold Spring Harbor Laboratory; 2019 Feb 26;4:557132.

30. Mondragon A, Davidsson D, Kyriakoudi S, Bertling A, Gomes-Faria R, Cohen P, et al. Divergent effects of liraglutide, exendin-4, and sitagliptin on Beta-cell mass and indicators of pancreatitis in a mouse model of hyperglycaemia. PLoS ONE. 2014;9(8):e104873.

31. Sisley S, Gutierrez-Aguilar R, Scott M, D’Alessio DA, Sandoval DA, Seeley RJ. Neuronal GLP1R mediates liraglutide’s anorectic but not glucose-lowering effect. J Clin Invest. 2014 Jun;124(6):2456–63.

32. Tølbøl KS, Kristiansen MN, Hansen HH, Veidal SS, Rigbolt KT, Gillum MP, et al. Metabolic and hepatic effects of liraglutide, obeticholic acid and elafibranor in diet-induced obese mouse models of biopsy-confirmed nonalcoholic steatohepatitis. World J Gastroenterol. Baishideng Publishing Group Inc; 2018 Jan 14;24(2):179–94.

33. Liang Y-L, Khoshouei M, Glukhova A, Furness SGB, Zhao P, Clydesdale L, et al. Phase-plate cryo-EM structure of a biased agonist-bound human GLP-1 receptor-Gs complex. Nature. Nature Publishing Group; 2018 Feb 21.

34. Lau J, Bloch P, Schäffer L, Pettersson I, Spetzler J, Kofoed J, et al. Discovery of the Once-Weekly Glucagon-Like Peptide-1 (GLP-1) Analogue Semaglutide. J Med Chem. 2015 Sep 24;58(18):7370–80.

35. Knudsen LB, Lau J. The Discovery and Development of Liraglutide and Semaglutide. Front Endocrinol (Lausanne). 2019;10:155.

36. Hupe-Sodmann K, McGregor GP, Bridenbaugh R, Göke R, Göke B, Thole H, et al. Characterisation of the processing by human neutral endopeptidase 24.11 of GLP-1(7-36) amide and comparison of the substrate specificity of the enzyme for other glucagon-like peptides. Regul Pept. 1995 Aug 22;58(3):149–56.

37. Zhu L, Almaça J, Dadi PK, Hong H, Sakamoto W, Rossi M, et al. β-arrestin-2 is an essential regulator of pancreatic β-cell function under physiological and pathophysiological conditions. Nat Commun. 2017 Feb 1;8:14295.

38. Thompson A, Kanamarlapudi V. Agonist-induced internalisation of the glucagon-like peptide-1 receptor is mediated by the Gαq pathway. Biochem Pharmacol. 2015 Jan 1;93(1):72–84.

39. Secher A, Jelsing J, Baquero AF, Hecksher-Sørensen J, Cowley MA, Dalboge LS, et al. The arcuate nucleus mediates GLP-1 receptor agonist liraglutide-dependent weight loss. J Clin Invest. 2014 Oct;124(10):4473–88.

40. Steen A, Larsen O, Thiele S, Rosenkilde MM. Biased and g protein-independent signaling of chemokine receptors. Front Immunol. 2014;5(1):277.

